# Probing the killing potency of tumor-infiltrating lymphocytes on microarrayed autologous tumoroids

**DOI:** 10.1101/2021.03.30.437679

**Authors:** Devanjali Dutta, François Rivest, L. Francisco Lorenzo-Martín, Nicolas Broguiere, Lucie Tillard, Simone Ragusa, Nathalie Brandenberg, Sylke Höhnel, Damien Saugy, Sylvie Rusakiewicz, Krisztian Homicsko, George Coukos, Matthias P. Lutolf

## Abstract

Immunotherapy has shown promise as an approach to fight cancer by harnessing the immune system. However, due to the lack of biomarkers to guide treatment regimens and predict response rates, there is an unmet need for more robust *ex vivo* and *in vitro* systems that recapitulate patient-specific tumor biology and enable response prediction for immune therapies in an autologous setting. To address this issue, we developed a high-content screening-compatible assay based on microcavity arrays to study tumor-infiltrating lymphocyte (TIL) functionality on 3D tumoroid models. We validated our system using the pmel-1 activated T cell mouse model to assess both cancer immunogenicity and T cell functionality. To demonstrate the translational potential of the platform, we used it to evaluate the response of patient-derived TILs to autologous human colorectal cancer (CRC) tumoroids. Using a combination of imaging and flow cytometry, we determined several features of the antitumor activity of TILs, including the extent of tumoroid killing and secretion of cytokines. We then used the approach to identify responders to immunotherapy, such as the immune checkpoint blockade (ICB) agent Nivolumab (PD-1 inhibitor) and Ipilimumab (CTLA-4 inhibitor). Our system allows not only the identification of immunogenic tumors, but also the testing of patients for response to immunomodulators, enabling personalized immuno-oncology.

## Introduction

Immunotherapy has become the fourth tier of cancer care, alongside surgery, chemotherapy and radiotherapy. After initial success in the treatment of melanoma^1^, it has quickly established itself as an important treatment modality for several types of solid cancers, including a subset of colorectal cancers (CRCs)^1^. Approved immunotherapy approaches used to treat cancer include immune checkpoint blockade (ICB), chimeric antigen receptor (CAR) T cell therapy, cytokines and oncolytic viruses^1,2^. Immune checkpoints are key modulators of the T-cell immune response that either promote or inhibit T-cell activation. The FDA-approved antibodies against the checkpoint T-cell receptor Programmed Cell Death Protein 1 (PD-1) have shown efficacy in patients with mismatch-repair-deficient and microsatellite instability-high (dMMR-MSI-H) CRC^2^. Blocking immune checkpoints is also efficient in malignant melanoma, nonsmall-cell lung cancer, renal cell cancers, hypermutated gastrointestinal cancers, and others^2^. However, the response rate of patients to immune checkpoint therapy with PD-1 blockade remains limited to 15-20% of the patients^2^. Moreover, it remains largely unclear why and which patients will respond to ICB^3, 4^. Therefore, assessing sensitivity to PD-1 blockade remains a challenge for ICB therapy in the clinic and requires the development of novel assay systems that have the potential to predict patient-specific responses.

Over the last decade, sophisticated three-dimensional (3D) *in vitro* tumor models have been developed^5,6,7^. These systems, often referred to as cancer organoids or tumoroids, mimic major aspects of the *in vivo* tumor biology and heterogeneity, including the production of extracellular matrix components, the distribution of oxygen and metabolites, the proliferation rates, and gene expression patterns^8,9^. Therefore, cancer organoids promise to advance the development of *in vitro* assays for testing immunotherapeutics and TIL functionality^8,10–12^. This is supported by the fact that melanoma cells grown as 3D spheroids exhibit more physiologically relevant immunomodulatory function than their 2D-cultured counterparts^13,14^. In addition, CRC cancer organoids are also better suited to detect patient response to CRC therapy^15,16^.

For an *in vitro* system to adequately recapitulate tumor biology, it should allow for interactions between tumor and its microenvironment through both cell-cell contacts and paracrine signaling factors such as cytokines, chemokines, and growth factors. In recent years, researchers have turned to organotypic spheroids, derived from both murine and human samples, to study changes in immune cell populations and response to immune checkpoint blockade using various 3D platforms^3,4,6^. Although the current 3D assay systems have enabled insightful studies, they are usually relatively low-throughput, do not allow easy and controlled co-culture establishment, and require high quantities of ill-defined extracellular matrix materials such as Matrigel. Bioengineered devices could mitigate some of these shortcomings and have the potential to facilitate personalized immuno-oncology efforts in a controlled and scalable setting.

Here, we sought to develop a reproducible and scalable 3D heterotypic tumoroid model to test TIL functionality and response to immunotherapeutic agents. To this end, we have created arrays of size-defined tumoroids in hydrogel-based microcavities providing up to 70 replicates per array on which lymphocytes can be easily added in various effector to target (E:T) ratios and in the presence or absence of immunomodulators^17^. Using 3D imaging and semiautomated image analysis, we quantified various features of heterotypic tumoroids, including cell death, tumoroid size, and TIL migration. In addition, lymphocyte phenotype and cytokine/chemokine secretion were determined by flow cytometry. The platform was first validated using a pmel-1 transgenic mouse model carrying altered T cell receptors to recognize the melanoma cell line B16-F10. This highlighted its potential to measure the tumor-killing potential of lymphocytes in a controlled and quantitative manner. To see if our approach could be useful in a preclinical setting, we then evaluated the functionality of patient-derived TILs on autologous CRC tumoroid derived from multiple patients. We demonstrate the ability of the system to distinguish patients with ‘cold’ or unresponsive tumors from those that are effectively targeted by T cells and/or respond to immune checkpoint blockade. Overall, these experiments demonstrate the potential of our platform for applications in precision medicine.

## Results

The Gri3D^™^ culture platform consists of hydrogel films, located at the bottom of wells in a 96-well plate, comprising arrays of round-bottom microcavities (or microwells) which allow for highly efficient cell aggregation in a solid matrix-free culture setting^18^. The microcavities can be generated with various sizes and numbers, e.g., 300 μm (~200 microcavities/well) and 500 μm (~70 microcavities/well). Unlike traditional organoids grown in 3D solid matrices, the microcavity array system promotes relatively homogeneous organoid growth in predefined positions, thereby allowing enhanced cell-cell interactions. It therefore allows reliable tracking and downstream processing of individual organoids and T cell activity in high-throughput. For our experiments we used the 96-well plates with 500 μm microcavities (**Figure 1a,b**).

**Figure 1:**
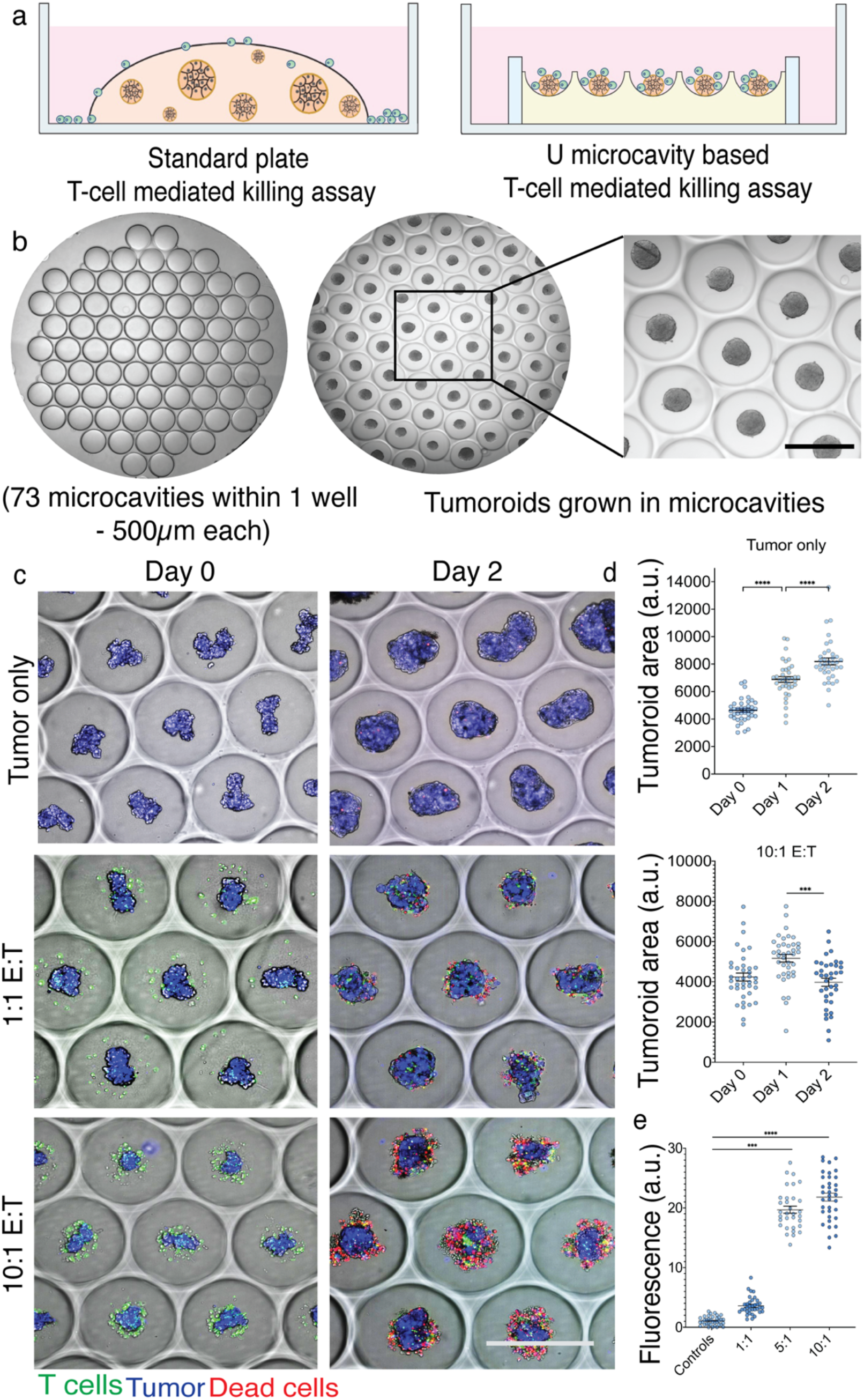
The microcavity array platform allows efficient assessment of T cell killing activity in the pmel-1 mouse model. (a) Illustration of the comparison between standard and microcavity-based culture systems in the setup of T cell killing assays. (b) Representative brightfield images showing the array of 500-μm microcavities used in this study before (left) and after (right) tumor cell seeding. Scale bar: 500μm. (c) Representative images of microcavities with individual tumoroids surrounded by T cells at the indicated effector to target (E:T) ratio (0:1; 1:1 or 10:1) and co-culture time (day 0 or 2). Tumor cells, T cells and dead cells were labeled with far-red cell tracer (blue), CFSE (green) and PI (red), respectively. Scale bar: 500μm. (d) Dot plots showing the tumoroid area over time at the indicated E:T ratios in the experiment shown in (c). ***, *P* < 0.001 (ANOVA and Tukey’s HSD test). (e) Measurement of the PI fluorescence intensity at day 2 in the indicated conditions from the experiment shown in (c). ***, *P* < 0.001 (ANOVA and Tukey’s HSD test).

To validate the suitability of this platform for the efficient detection and quantitation of T cell anti-tumor activity, we made use of the pmel-1 mouse model. These animals bear T cells with a transgenic TCR receptor able to recognize the gp100 antigen expressed by murine B16 melanomas, ensuring immune cell responsiveness and tumor cell killing when these two cell types are co-cultured. We seeded B16-F10 melanoma cells in the microcavity system, allowed tumoroid formation for two days, and then added freshly extracted pmel-1 T cells at different effector-to-target ratios (**Figure 1c**). We were able to monitor at the single-organoid level how T cells migrated towards cancer cells and exerted their cytotoxic activity, which led to the disruption of tumoroids over time (**Figure 1c,d**). In contrast, melanoma tumoroids cultured in the absence of T cells steadily grew in size (**Figure 1c,d**). To quantify T-cell killing efficacy, we stained dead cells by adding propidium iodide (PI) to the microcavities (**Figure 1c**). We found a direct correlation between the killing efficacy and the effector-to-target ratios, showing the sensitivity of the system to the amount of T cells (**Figure 1e**).

We next evaluated the translational potential of this platform by focusing on the probing of T-cell activity in colorectal cancer, where immunotherapy is applied according to tumorspecific markers^4^. Cancer cells and tumor-infiltrating lymphocytes were isolated from patient-derived CRC biopsies, expanded *in vitro* (**Figure 2a**), and also mutationally characterized by targeted Next-Generation Sequencing of tumoroid genomic DNA (unpublished data). We generated five different patient-derived tumoroid lines, two of them carrying mutations in key DNA repair genes (e.g., *ATM* in patient #3, *MSH6* and *NBN* in patient #4), a distinctive feature of ICB-responsive CRC tumors^2^.

**Figure 2:**
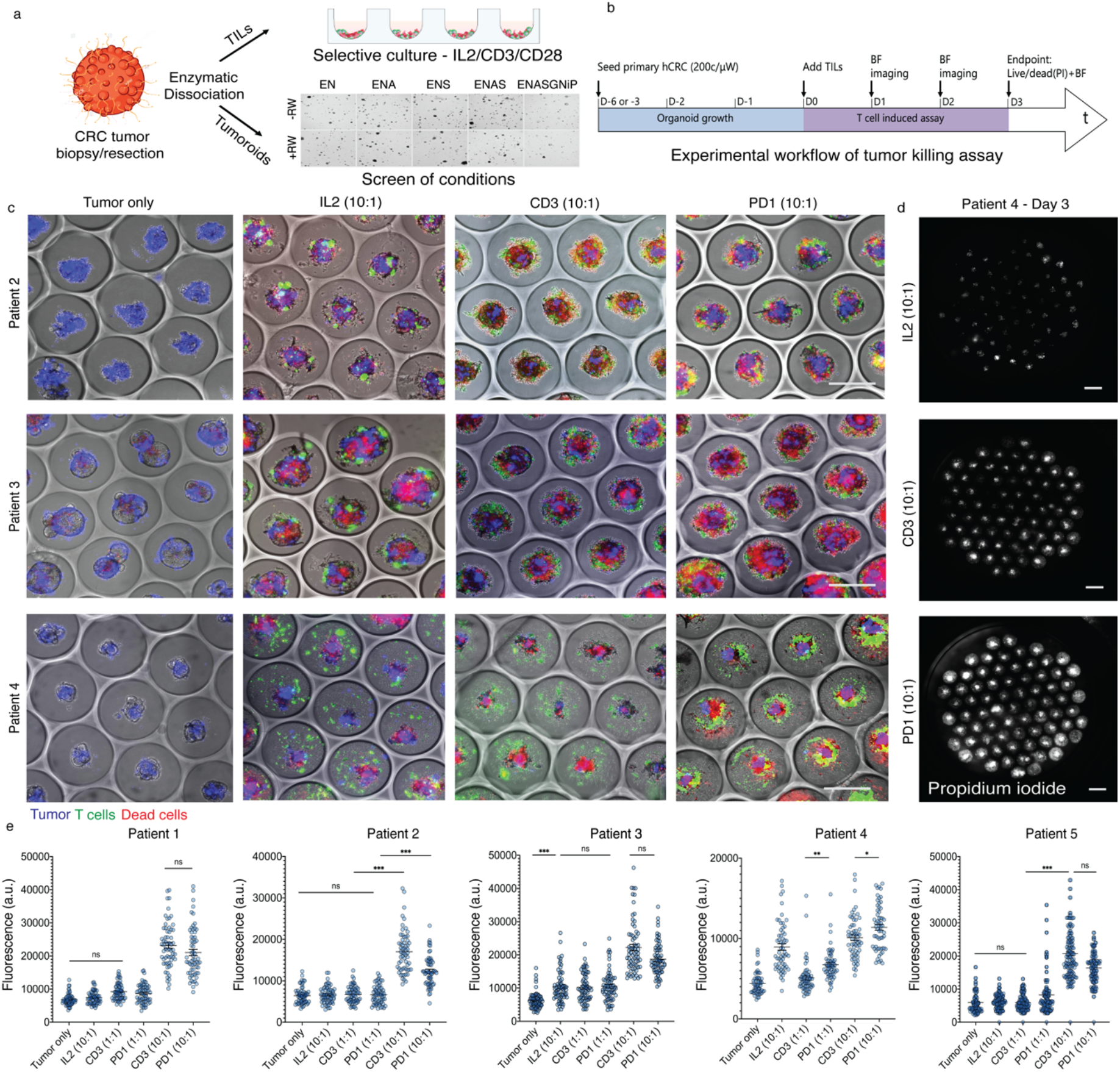
The microcavity array platform allows screening of TIL activity in human colorectal cancer samples. (a) Illustration of the protocol followed in this study to isolate tumor-infiltrating lymphocytes and derive tumoroids from CRC samples. (b) Scheme of the experimental workflow followed in this study to perform T cell killing assays (c) Representative images of microcavities with individual tumoroids surrounded by T cells in the indicated conditions and patients on day 3. Tumor cells, T cells and dead cells were labeled with far-red cell tracer (blue), CFSE (green) and PI (red), respectively. Scale bar: 500μm. (d) Representative images showing propidium iodide staining in microcavities seeded with cell from patient #4 in the indicated conditions at day 3. Scale bar: 500μm. (e) Quantification of PI staining intensity in the indicated patients and conditions from the experiment shown in (c). ns, *P* > 0.05; *, *P* < 0.05; **, *P* < 0.01; ***, *P* < 0.001 (ANOVA and Tukey’s HSD test). Culture conditions: IL2 10:1 (IL2, 10:1 E:T); CD3 1:1 (IL2 + α-CD3/CD28; 1:1 E:T), CD3 10:1 (IL2 + α-CD3/CD28, 10:1 E:T); PD1 1:1 (IL2 + α-CD3/CD28 + α-PD1/α-CTLA4, 1:1 E:T); PD1 10:1 (IL2 + α-CD3/CD28 + α-PD1/α-CTLA4, 10:1 E:T).

Co-cultures in the microcavities were established according to the workflow indicated in **Figure 2b**. As a positive control for T cell responsiveness, we co-stimulated TCR signaling ectopically with α-CD3 and α-CD28 antibodies (**Figure 2c,e**). Under these conditions, all tumoroid lines tested were capable of eliciting a cytotoxic response in at least one of the effector-to-target ratios tested (**Figure 2c,e; Figure 3a**). Using conditions deprived of α-CD3 and α-CD28, we found that the TILs from two patients (#3 and #4) were able to autonomously recognize and target the autologous cancer cells (**Figure 2c, e; Figure 3a; Suppl. Figure 1c**). This correlated with an increased capability of the TILs to migrate towards the tumoroid (**Suppl. Figure 1a,b**).

**Figure 3:**
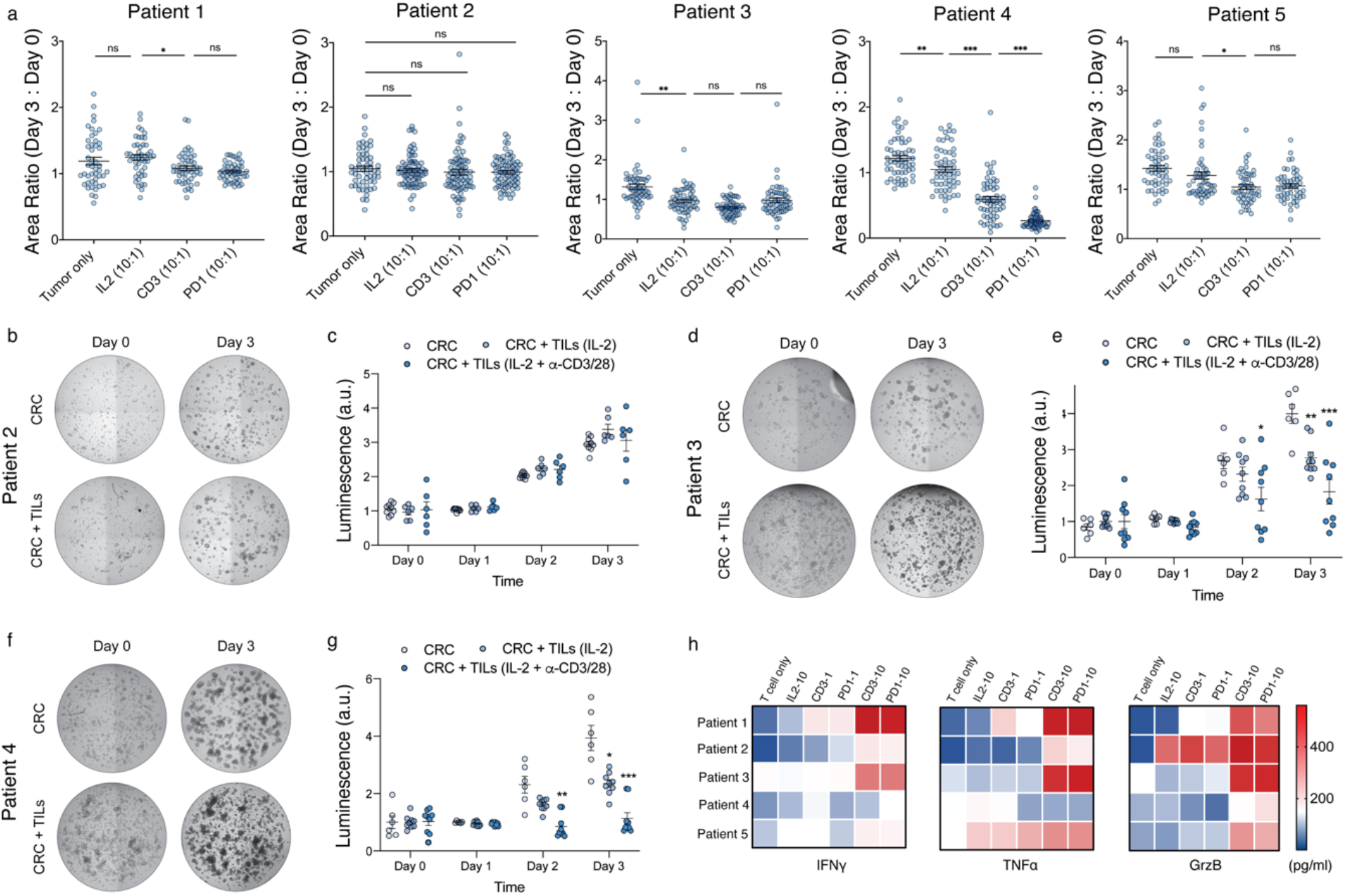
Luciferase-reporter assays and cytokine profiling support the readouts from the microcavity system. (a) Quantification of tumoroid area over time for the indicated patients and conditions. *, *P* < 0.05; **, *P* < 0.01; ***, *P* < 0.001 (ANOVA and Tukey’s HSD test). (b-g) Representative brightfield images (b, d, f) and luminescence quantifications (c, e, g) of the indicated co-cultures of luciferase-labeled CRC organoids and autologous TILs. *, *P* < 0.05; **, *P* < 0.01; ***, *P* < 0.001 (ANOVA and Dunnett’s multiple comparisons test). (h) Heatmaps showing the amount of granzyme B (GrzB), interferon γ (IFNγ) and tumor necrosis factor α (TNFα) released by the TILs of the indicated patients in the indicated conditions. Culture conditions: IL2 10:1 (IL2, 10:1 E:T); CD3 1:1 (IL2 + α-CD3/CD28, 1:1 E:T); CD3 10:1 (IL2 + α-CD3/CD28, 10:1 E:T); PD1 1:1 (IL2 + α-CD3/CD28 + α-PD1/α-CTLA4, 1:1 E:T); PD1 10:1 (IL2 + α-CD3/CD28 + α-PD1/α-CTLA4, 10:1 E:T).

Next, to assess response to immunotherapeutic agents currently used in the clinic, we included the immune checkpoint blockade antibodies Nivolumab (PD-1 inhibitor) and Ipilimumab (CTLA-4 inhibitor) in the assay (**Figure 2c,e**). While this did not translate into increased cytotoxicity in most patients, T-cell activity was significantly increased in co-cultures from patient #4 (**Figure 2c-e**; **Figure 3a**; **Suppl. Figure 1d**). Taken together, these results highlight the adaptability of our microcavity array-based approach for assessing T-cell responsiveness in different immunomodulatory contexts (**Suppl. Figure 2**).

To validate the specificity of the microcavity system, we labeled our patient-derived tumoroids with a luciferase reporter that allows precise quantification of viability. These tumoroids were cultured with TILs in Matrigel ‘drops’ on standard 96-well plates (**Figure 1a**, left scheme) under the same conditions used for the microcavity arrays. By measuring CRC luminescence on consecutive days, we confirmed that the TILs derived from patients #3 and #4 were capable of affecting tumor cell growth in an autonomous manner (**Figure 3d-g**, IL-2 condition). This anti-tumor activity was enhanced upon α-CD3/CD28 co-stimulation (**Figure 3d-g**, IL-2 + α-CD3/CD28 condition). Conversely, we did not detect CRC growth impairment by the TILs isolated from patient #2 (**Figure 3b,c**). Therefore, results obtained from the microcavity system can be reproduced in a standard low throughput culture model.

Finally, we tested whether the cytotoxicity observed in the microcavity setup was associated with other features of T-cell activation, such as cytokine production. To this end, we profiled the release of the cytotoxic factors Granzyme B (GrzB), Interferon-γ (IFNγ), and Tumor Necrosis Factor-α (TNFα) in the supernatant of the microcavities. As a positive control, we found that both higher effector-to-target ratios and co-stimulation with α-CD3/CD28 promoted cytokine production (**Figure 3h**). This response was enhanced by ICB agents only in the case of TILs derived from patient #4, in line with our previous results (**Figure 3h**; **Figure 2**). However, the level of cytokine production did not correlate with cytotoxicity in all TIL lines (**Figure 3h**), suggesting that the mechanism mediating anti-tumor activity of TIL may depend on additional patient-specific factors (**Suppl. Figure 2**).

## Discussion

The complex and heterogeneous nature of tumors makes personalized treatment a prerequisite for most cancer therapies to achieve optimal response rates. This is particularly true for immunotherapy, where patient-specific immunological factors further increase the complexity of drug responses^1–4^. Although the presence of certain molecular markers, such as a high mutational burden, can be helpful in predicting response to immunotherapy, platforms that can functionally validate such predictions in a reliable and efficient manner are still needed. Here, we demonstrate that our microcavity-based tumoroid culture system can fill this gap by providing a unique experimental setup in which a large number of uniform tumoroids can be robustly examined at the single-tumoroid level using both microscopy and flow cytometry workflows. Unlike other recently developed co-culture systems that require the use of complex extracellular matrices or do not allow precise control over cell distribution and organoid/spheroid size^5–7,12^, our platform provides unparalleled resolution of tumoroid-TIL interactions in a minimal, matrix-free setup.

Using this system, which demonstrates its potential for clinical translation, we were able to stratify colorectal cancer patients according to both tumor-infiltrating lymphocyte reactivity and response to immune checkpoint blockade. Although attribution of these results to clinical outcomes and expansion of the study to a larger number of patients to assess the translational reliability of the model are ongoing, our results are consistent with current clinical evidence for immunotherapy in colorectal cancer^4^. Specifically, the microcavity setup demonstrated increased immunogenicity in patients with mutations in key genes involved in DNA repair. In addition, the patient who responded to ICB (patient #4) carried a mutation in *MSH6*, a marker of mismatch repair deficiency and immunotherapy benefit in CRC^4^. Taken together, these results support the suitability of the microcavity platform as a novel tool for personalized immunotherapy testing.

## Materials and Methods

### Pmel-1 T-cells B16-F10 Mouse killing assay

#### B16-F10 staining and plating

B16-F10 melanoma cells were received as a kind gift from the Laboratory of Biomaterials for Immuno-engineering (LBI EPFL). First, the cells were stained with a far-red cell tracer (CellTrace™ Far Red Cell Proliferation Kit; Molecular Probes) as follows: cells were resuspended in 1ml warm PBS with tracer at 1:1000 and incubated 20min at 37°C. Then equal amount of full medium with 10% FBS was added and incubated 5 more minutes at 37°C. Suspension was centrifuged at 200g for 5 minute and cells were resuspended in 1ml of full medium (DMEM + 10% FBS; Gibco). Cells were either plated in 2D on cell culture treated flat bottom 96-well plates (Falcon) at 3’000 cells per well and supplemented with 150μl full medium, or in 3D in 96-well Gri3D^™^ culture platform (Gri3D-96-S-24P; SUN bioscience) at 10K cells/per well, *i.e*., about 150 cells/microcavity. For 3D growth, the hydrogel microcavities were first equilibrated with full medium for 30min, then the medium was aspirated and a 20μl drop of cells at 10M cells/ml was carefully pipetted onto each microcavity array. The cells were let to sediment for 30min, before an additional 150μl of cold full medium containing 2% Matrigel was added slowly in the side well. Adding 2% Matrigel was necessary for the proper formation of the cysts, by providing necessary ECM components. In both 2D and 3D, cells were incubated for 2 days before the T-cells co-culture with T-cells.

#### Pmel-1 mouse T-cells staining

Pre-activated Pmel-1 TCR transgenic T-cells were provided by the LBI (EPFL), suspended in culture medium containing IL-2 (10ng/ml; BioLegend). Cells were directly stained with CFSE (Cayman Chemical) and plated after reception. Briefly, cells were resuspended in a CFSE staining solution (CFSE stock solution diluted in assay buffer 1:200, followed by a dilution 1:2 in warm PBS) and incubated at 37°C for 20 minutes. Equal volume of medium containing 10% FBS was added and the solution was incubated 5 more minutes before centrifugation at 200g for 5 minutes. Cells were then resuspended in medium at the desired concentration, typically 10M cells/ml, and were ready for plating.

#### B16-F10 antigen spiking and Pmel-1 T-cells co-culture

To increase the response and cytolytic activity of the T-cells, the melanoma cells were spiked with gp100^+^ antigens (GenScript) just before addition of T-cells. Medium was aspirated from the wells for 2D and 3D cultures, and 100μl of medium containing the antigen at 1μM was added and incubated for 1h at 37°C. The spiked medium was then removed and the T-cells were added at different effector to target E:T ratios, either 1:1, 1:5, or 1:10, in function the initial cancer cells seeding number. In 2D, cells were added in 150μl of fresh T-cells medium (RPMI 1640 + 10% FBS + 10ng/ml IL2). In 3D, cells were plated carefully as 20ul drops on the microcavities array and let to sediment for 30 minutes, before 150μl of fresh T-cells medium was added in the side well.

#### Pmel-1 killing assay imaging

Images were taken 2 hours after co-seeding, and after 1 and 2 days of co-culture on an *INCell Analyzer 2200* (GE Healthcare), with a large field-of-view sCMOS camera (resolution 2048×2048 pixels, 16bits) and a 4x objective (0.2 NA). Wells were imaged in brightfield, far-red (645/705, tumoroids) and green channels (490/525, T-cells), with a software autofocus on the far-red channel. Images were taken as deconvoluted 3D stacks with flat field correction.

#### Image processing

Fiji software was then used to process the images and create maximal intensity projections of the 3D stacks. Brightfield images were used to define each microcavity of interest, within each well. When focus and contrast were good on the overall well, an automatic detection of the microcavities was possible by applying an *intermode* threshold (Supplemental Figure 3), followed by binary processing (*dilate, close, fill holes*) to represent each microcavity as a black circle. *Analyze particles* function was then used to detect every circle and store them as region of interest (ROI). When the automatic detection was not possible, black circles (radius of 250μm) were added manually on the microcavity of interest, before processing with the same thresholding and *analyze particles* function. Microcavities containing no cyst, more than one, or large debris were not included in the analysis. Once each microcavity ROI was defined, it was possible to analyze them individually.

#### Measuring tumoroid area

To measure cysts area within each microcavity, the far-red channel was used to set a threshold (*triangle*). When the tumoroids fluorescence was homogenous, it was possible to use the *analyze particle* function to measure their area and shape. However, in the majority of the cases, the tumoroids intensity faded over time in co-culture and only an estimation of the area could be done by measuring the overall mean fluorescence intensity in the whole microcavity after applying the threshold.

#### Measuring cell death by PI intensity

PI intensity was measured in an area of interest 50μm larger than the cysts. To define the extended perimeter, the tumoroids were thresholded as before, and the binary function *dilate* was used (60 times, count at 3). *Analyze particle* function was then necessary to detect each area and save them as ROI. Before measuring PI intensity, a compensation was required due to the spectral overlap with the green fluorescence. First, a background subtraction (rolling ball radius of 50px) was performed on both fluorescent channels, then the *image calculator* function was used to subtract the green intensity (CFSE) from the red channel (PI). The resulting compensated image was used to measure the PI mean intensity within each area of interest.

#### Measuring cell death by flow cytometry

Cells were recovered from the microcavity array by pipetting up and down several times 100ul of medium, and transferred into FACS tubes (Falcon) for dissociation and staining. The tubes were then centrifuged at 200g for 4min, and cells resuspended into warm 200ul Trypsin (Life Technologies) for 10 minutes. 1ml of medium containing 10% FBS was added into each tube to block the enzymatic process. Each sample was then strained through FACS tubes strainer cap (5ml, 35μm strainer, Corning) and washed twice with PBS to recover most cells. The cells were then centrifuged at 200g for 4 minutes, washed once in 200μl of PBS, and resuspended in 50μl staining solution. A simple live-dead staining was done with a LIVE/DEAD™ Fixable Aqua kit (1:1000; Molecular Probes) for 30 minutes in the fridge. The samples were washed twice with PBS, resuspended in 300ul PBS, and analyzed on an *LSRFortessa* flow cytometer (BD Biosciences).

### Isolation and establishment of CRC tumoroids

Tumor sample was washed 1x in a 50 ml tube with Advanced basal DMEM. Cells were maintained on ice. Tumor samples were minced into small pieces in 2-3 ml media with a surgical blade. The pieces were collected with forceps in a new 50ml tube and digested in 10ml BM with: Collagenase I [0.1g/ml], Hyaluronidase [20mg/ml], BM = Advanced DMEM/F12 +1% P/S +Glutamine +HEPES. Digest for 30’ at 37°C in the water bath. Cells were filtered with a 70 μm strainer in a new 50 ml tube (Phase 1). 1ml FBS was added to the digested solute to block the reaction. Remaining tissue pieces were collected in another 50 ml tube by inverting the strainer and washing it with 10 ml Tryple 1x solution. The tissue pieces were dissociated in Tryple 10’ at 37°C with agitation. The pieces were collected again with a strainer, in a new tube and, blocked with 1 ml FBS (Phase 2). Phase 1 and Phase 2 were pooled together as: First Digest (FD). An FD fraction was plated in flat and round bottom plates ±stimulation in T cells medium (recipe below) Another FD fraction was used for the tumor cells, and was plated in Matrigel in 24 well plates, ±hypoxia. Ideally 10 drops, 20 μl each were plated into 10 Conditions in DMGF (recipe below): EN, ENA, ENS, ENAS, ENASgNiP, EN + RW, ENA +RW, ENS +RW, ENAS +RW, ENASgNiP + RW. The culture media used was 500μl/well. Other components i.e. Primocin 1:500, Thiazovivin 1:5000, R-Spondin 1.5:1000 are added in each well. E: EGF, N: Noggin, A: A83-01 (TGFβi), S: SB202190 (p38i), W: Wnt3A, R: R-spondin, G: Gastrin, Ni: Nicotinamide, P: Prostaglandin-E2.

*T cell media (for 500 ml)* = RPMI 1640 + Glutamax (1X) (432ml) + FBS 10% (50ml) + 2-mercaptoethanol 50nM (500ul) + P/S (5ml) + HEPES 1 M (12.5ml) (25mM) *DMGF (for 50ml)* = BM+B27 (1 ml) + N-acetyl cysteine (NAC, 100 ul) (Stable 30 days).

### Exome sequencing and analysis

Organoids were collected in DMEM to remove matrigel and subsequently washed twice in PBS. Genomic DNA was isolated using the PureLink^™^ Genomic DNA Mini Kit (ThermoFisher, Catalog No. K182002) as per the manufacturer’s instructions. Exome sequencing was performed by BGI Genomics (Hong Kong) at 100X depth using 150 bp paired-end reads. Sequenced reads were mapped to human genome assembly GRCh38 using BWA-MEM (v0.7.17). For variant calling, Samtools (v1.9) was used to detect substitutions and indels. For variant annotation, we used the Ensembl Variant Effect Predictor (v100.2).

### Culture and expansion of TILs

Using an ultra-low adherence 96 WP with round bottom well allows for rapid evaluation of the TILs proliferation and great for comparing culture conditions. Cells from the tumor dissociation were suspended in T cell media containing RPMI 1640, FBS, HEPES, Pen strep and 2-mercaptoethanol (See recipe above). Recommended numbers of cells for suspension: 10K cells in 200ul volume per well of a 96-UWP, and 50K per well in 400ul of a 48WP. Media was replenished every 2 days. Recommended volume to remove: 90ul for the 96uWP, 240ul for the 48WP. Fresh medium with IL2 was added at 2x concentration (6000U/ml), 100ul for the 96uWP and 250ul for the 48WP. TILs were expanded by dissociating the aggregates and expanding in suspension cultures in the ratio 1:2 or 1:3.

### Aggregating tumor cells in microcavities

Gri3D plates (500um) were obtained from Sun Biosciences. The plate can be kept in the fridge for few weeks in PBS, and usable for more than one experiment. The desired hydrogel microcavities were equilibrated with medium (avoid most outer wells as they dry faster, instead keep them filled with PBS to prevent edge effect). PBS was aspirated and 150ul of DMGF per well was added and let incubate (at least 30min). Matrigel drops were disrupted with 500ul cold TrypLE per well, mechanically breaking the drop by pipetting up and down. All cells were pooled in a 15ml falcon and placed in the water bath 37° for 8 min. 8ml of medium containing 10% FBS was added to block the TrypLE. Cells were centrifuged for 4min at 200g and resuspended in 500ul DMGF. Cells were stained with a tracer (for instance far red CellTrace) for 20min at 37°C. 2ml medium with FBS 10% was added to block. Cell were then centrifuged for 4min at 200g and resuspended in desired amount of complete medium (containing Thiazovivin) at a concentration of 20K-30K cells/ml. Medium was aspirated from the desired microwells using the notch on the wall and cells were seeded in the microcavities in 20ul to 40ul drops for an equal sedimentation. Drops were allowed too sediment for a~30min, 150ul of cold complete medium containing T and 2% matrigel was added on top. The plate was maintained in the incubator and tumoroids were grown for 3 days.

### Co-culture establishment of tumoroids and T cells

#### TIL activation

If TILs are frozen, they require an overnight (to a full day) resting phase before stimulation to prevent cell death. TILs were thawed and plated in full RPMI medium with IL-2 at 100U/ml (resting medium). After 24h hour, they were activated with IL-2 at 3000U/ml and α-CD3/CD28 at 1ug/ml each for 1 day.

#### TIL Co-Seeding

Cells were stained with a tracer (for instance CFSE). CFSE was diluted in buffer 1:200. And cells were resuspended in the staining solution. Cell were incubated for 20min at 37°C. 2ml medium with FBS 10% was added to block. Cell were then centrifuge 4min at 200g. and resuspended in desired amount of full RPMI medium for a good concentration for the highest E:T ratio (e.g. 10^6 cells/ml). Seeding drop should be about 20 to 30ul. Medium from the microcavity containing tumoroids was aspirated from the side chamber and then the main chamber. Drop of TILs was added as centered as possible and allowed to sediment for 30min. 150ul of medium of choice with various factors/drugs were then added from the side chamber. PD-1 inhibitor was added at a concentration of 10mg/ml and CTLA-4 inhibitor was added at a concentration of 5mg/ml.

### T cell migration measurement

T cell migration was measured by quantifying the movement of T cells in conditions from Day 0 to Day 3. Spread of T cells around tumoroids was measured by computing the area of spread around the tumoroids and the radius of T cells around all tumoroids within a well on Day 0 and Day 3. A ratio of Day 0: Day3 indicated the degree of movement of T cells.

### Tumor killing assessment by measurement of PI and tumor shrinkage

To measure tumor killing by TILs, Propidium iodide was added to each well and imaged using InCell Analyzer. Imaging was performed with Z-stacks and in 4 quadrants which were then stitched in ImageJ. To measure tumor size, each tumoroid area was measured on Day 0 and Day 3 and ratio was computed. A ratio of >1 indicated tumor growth and <1 indicated tumor shrinkage over time.

### Cytokine analysis by CBA method

After the 2-3 days of co-culture, transfer 50ul of the supernatant from each microcavity in a round bottom 96WP and follow the CBA protocol for plates. Analysis was done in duplicates of each microcavity as there is about 130ul of supernatant available. Set up the instrument following the BD protocol (CBA setup) and analysis was done using the HTS cytometer module for 96WP format.

### Luciferase-based quantification of T cell killing activity

Lentiviral particles carrying a luciferase reporter were produced at the EPFL Gene Therapy Facility by transfecting HEK293 cells with the pHIV-Luc-ZsGreen plasmid (Addgene, Catalog No. 39196). Lentivirus-containing supernatants were collected and concentrated by centrifugation (1,500 g for 1 hr at 4°C). Lentiviral titration was performed using a p24-antigen ELISA (ZeptoMetrix, Catalog No. 0801111).

For lentiviral transduction, CRC organoids (~2×10^5^ cells) were collected in basal medium and dissociated into single cells by incubating in TrypLE (Thermo Fisher Scientific, Catalog No. 12605028) at 37°C for 5 min. CRC cells were then washed with basal medium 10% fetal bovine serum (Thermo Fisher Scientific, Catalog No. 10500064), resuspended in BMGF medium containing ~1000 ng(p24)/mL of lentiviral particles plus 8 μg/mL polybrene (Sigma-Aldrich, Catalog No. TR-1003-G), and plated in a 24-well plate. This plate was then centrifuged at 600 g for 60 min at room temperature and incubated for 6 hours at 37°C. After incubation, cells were collected, spun down, plated in 20 μL Matrigel drops (Corning, Catalog No. 354263) in a 24-well plate and cultured in BMGF medium. One week after infection, successfully transduced CRC cells (ZsGreen^+^) were sorted in a FACSAria Fusion flow cytometer (BD Biosciences).

For the luciferase-based monitorization of CRC cell viability, transduced CRC organoids (~10^4^ cells) were co-cultured with T cells (effector-to-target ratio of 10-to-1) in 8 μL Matrigel drops in a black-wall 96-well plate (Corning, Catalog No. CLS3603-48EA) and cultured in T cell medium. Where indicated, T cells were activated using 3000 U/mL IL-2 (PreproTech, Catalog No. 200-02-500), 1 μg/mL anti-CD3 antibody (BioLegend, Catalog No. 317326), and 1 μg/mL anti-CD28 antibody (BioLegend, Catalog No. 302934). Live-cell luciferase readings were performed for 3 consecutive days by adding 150 μg/mL luciferine (GOLDBIO, Catalog No. LUCK-500) and measuring luminescence in a Tecan Infinite F500 reader (Tecan).

## Acknowledgements

We thank L. Tang and K. Lei for providing cells for the mouse assay validations and S. Li for help with CRC sample handling, F. Kuttler and C. Claus for scientific discussions and inputs on the manuscript, and M. Gueye for help with cell culture. We acknowledge support from the following EPFL core facilities: Bioimaging and Optics, Flow Cytometry and Biomolecular Screening. This work was funded by the Personalized Health and Related Technologies (PHRT) Initiative from the ETH Board; and EPFL.

## Author contribution

M.P.L. and F.R. conceived the initial ideas. D.D., F.R. and M.P.L. designed experiments, analyzed data and wrote the manuscript. F.R. performed all experiments related to the mouse assay and conducted proof-of-concept experiments on patient-derived CRC tumoroids. D.D. performed and analyzed experiments on patient-derived CRC tumoroids. L.F.L.M. helped with experiments on human CRC samples, designed, performed and analyzed the luciferase-based experiments and helped in manuscript writing. N.B. spearheaded the establishment of the tumoroid/TIL biobank and cytokine assay analysis. S.R. (Ragusa) helped with biobank establishment and tumoroid characterization. N.B. and S.H. developed the microcavity platform and provided scientific inputs. L.T. helped with cytokine assay. S.R. (Rusakiewicz), D.S. and G.C. established the T cell protocols and provided critical immunology expertise. K.H. provided human CRC samples and information on the mutational profiles, contributed to experimental design and provided inputs on the manuscript. G.C. contributed to experimental design and provided inputs on the manuscript.

---

**Supplementary Figure 1:**
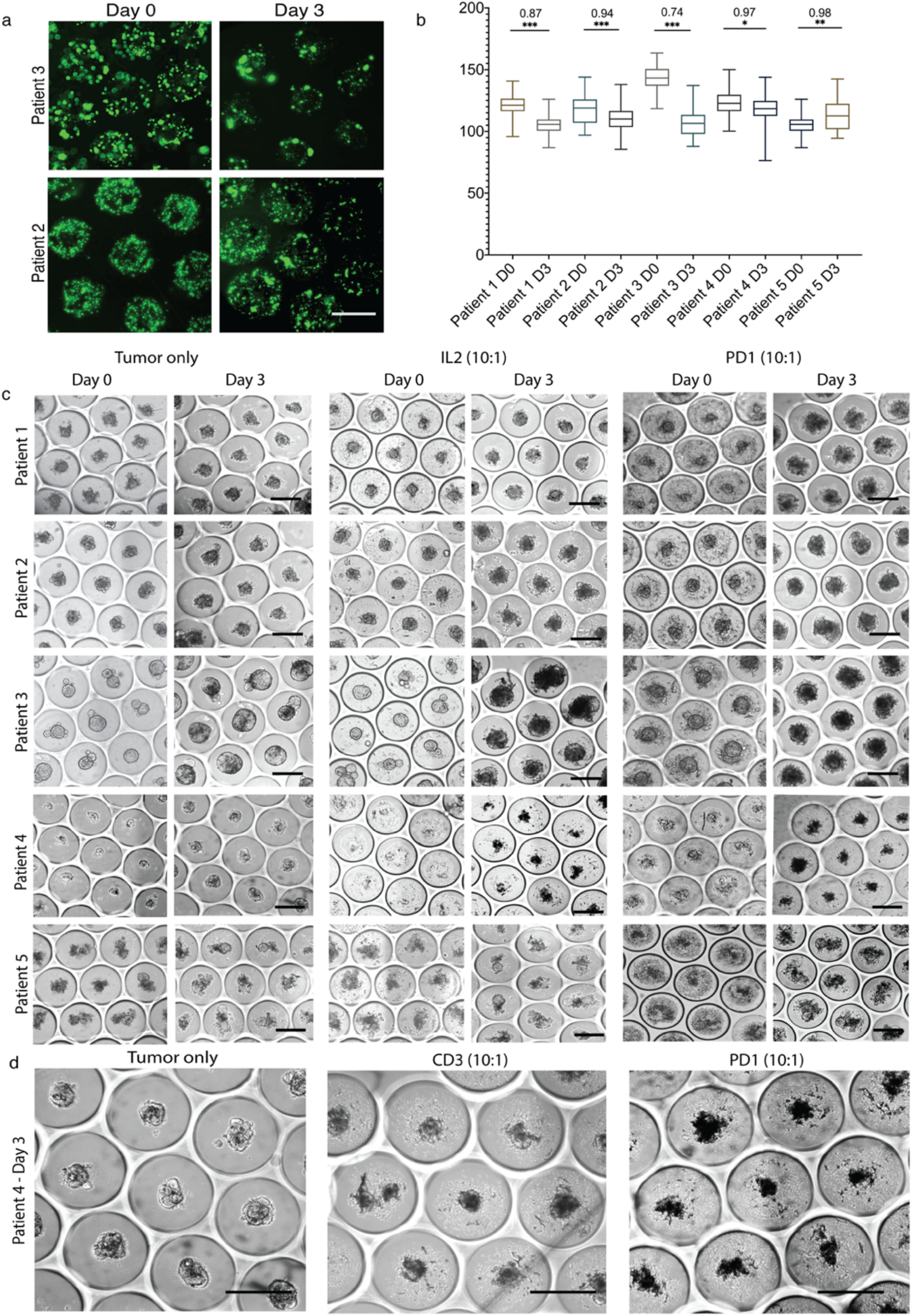
Microcavity-based T cell killing assay allows visual inspection of cytotoxic activity. (a) Representative example images illustrating TIL migration during the co-culture with autologous CRC for the indicated patients. TILs are labeled in green. Scale bar: 500μm. (b) Quantification of TIL migration between day 0 and day 3 for the indicated patients in IL2 only condition. Values indicate the ratio of radius between Day 3 and Day 0. *, *P* < 0.05; **, *P* < 0.01; ***, *P* < 0.001 (*t*-test). (c) Representative images of microcavities with individual tumoroids surrounded by T cells in the indicated conditions, patients and days. Scale bar: 500μm. (d) Representative images of microcavities with individual tumoroids surrounded by T cells in the indicated conditions for patient #3 on day 3. Scale bar: 500μm. Culture conditions: IL2 10:1 (IL2, 10:1 E:T); CD3 1:1 (IL2 + α-CD3/CD28, 1:1 E:T); CD3 10:1 (IL2 + α-CD3/CD28, 10:1 E:T); PD1 1:1 (IL2 + α-CD3/CD28 + α-PD1/α-CTLA4, 1:1 E:T); PD1 10:1 (IL2 + α-CD3/CD28 + α-PD1/α-CTLA4, 10:1 E:T).

**Supplementary Figure 2:**
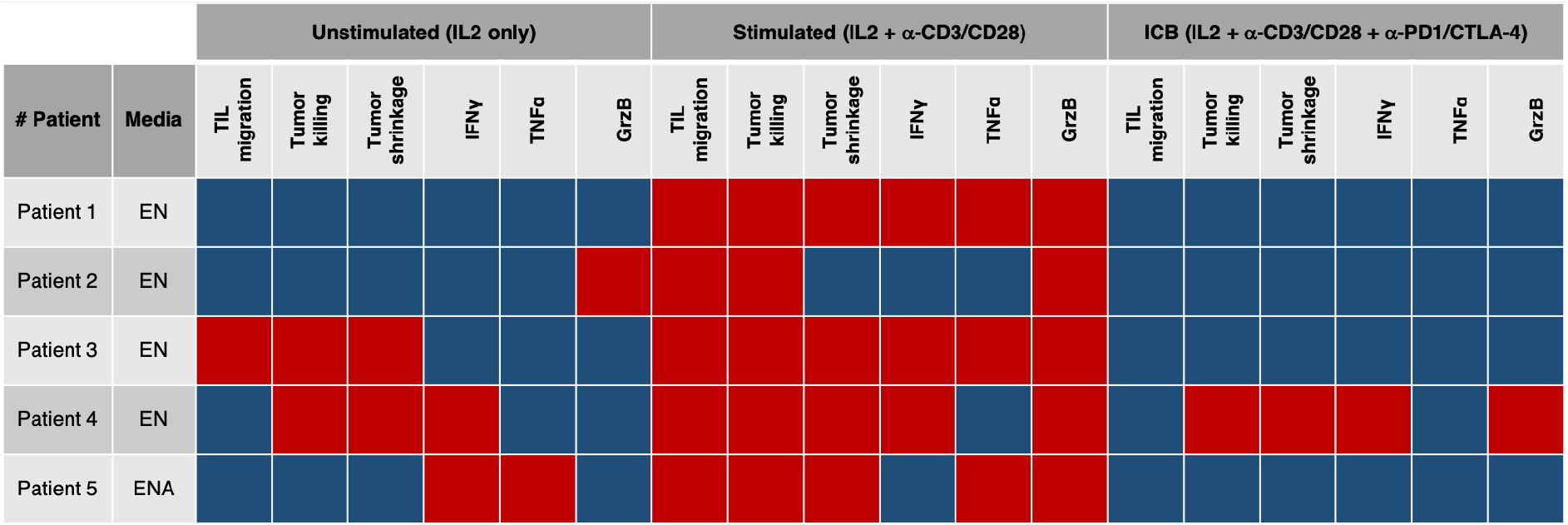
Summary of patient responses in microcavity-based T cell killing assay. Table shows a qualitative global comparison of responses observed in different assays. Shown are the growth factors added in basal media for expansion of tumoroids of each patient. Red: represents a significant effect or increase. Blue: represents no effect or no increase. For comparison, unstimulated conditions were compared to tumor only conditions such that in red are conditions where there was an increase in the presence of native T cells. Stimulated conditions were compared to unstimulated conditions, such that red shows an increase due to the activity of α-CD3/CD28. ICB conditions were compared to stimulated conditions such that red shows increase due to the activity of immune checkpoint inhibitors: α-PD1/α-CTLA4. Culture conditions: Non-stimulated (IL2, 10:1 E:T); stimulated CD3 10:1 (IL2 + α-CD3/CD28, 10:1 E:T); and ICB PD1 10:1 (IL2 + α-CD3/CD28 + α-PD1/α-CTLA4, 10:1 E:T).

